# An Open-Source Computational Tool for Measuring Bacterial Biofilm Morphology and Growth Kinetics upon One-Sided Exposure to an Anti-Microbial Source

**DOI:** 10.1101/2022.04.20.488818

**Authors:** Sarah Gingichashvili, Doron Steinberg, Ronit V. Sionov, Osnat Feuerstein, Noa E. Cohen

## Abstract

*Bacillus subtilis* biofilms are well known for their complex and highly adaptive morphology. Indeed, their phenotypical diversity and intra-biofilm heterogeneity make this gram-positive bacterium the subject of many scientific papers on the structure of biofilms. The ‘robustness’ of biofilms is a term often used to describe their level of susceptibility to anti-microbial agents and various mechanical and molecular inhibition/eradication methods. In this paper, we use computational analytics to quantify *Bacillus subtilis* morphological response to proximity to an anti-microbial source, in the form of the antiseptic chlorhexidine. Chlorhexidine droplets, placed in proximity to *Bacillus subtilis* macrocolonies at different distances result in morphological changes, quantified using Python-based code, which we have made publicly available. Our results quantify peripheral and inner core deformation as well as differences in cellular viability of the two regions. The results reveal that the inner core, which is often characterized by the presence of wrinkled formations in the macrocolony, is more preserved than the periphery. Furthermore, the paper describes a crescent shaped colony morphology which occurs when the distance from the chlorhexidine source is 0.5 cm, as well as changes observed in the growth substrate of macrocolonies exposed to chlorhexidine.

## Introduction

Biofilms are often described in terms of their ‘robustness’ – the term embodies a set of characteristics that allow for bacteria to thrive within the biofilm. The set of features that makes up the ‘robustness’ of biofilms is diverse – biofilm height, cellular differentiation, strength of adherence to substrate and surface hydrophobicity have all been previously used to assess key biofilm features^1–3^. Such ‘robustness features’ can roughly be characterized into features that relate to either macrocolony structure (e.g., biofilm height and internal structure), cellular composition (e.g., amount of secreted exopolysaccharides (EPS), quorum-sensing, surface hydrophobicity) or effectiveness of response to stress conditions (e.g., nutrient limitation, anti-bacterial agents).

*Bacillus subtilis* (*B. subtilis*) is a model organism that has traditionally been studied for its complex biofilm macrocolonies^4^. Mature *B. subtilis* macrocolonies are characterized by a well-defined central core that differs morphologically from colony periphery. Furthermore, *B. subtilis* biofilms have been found to be highly adaptive in terms of their morphology to changing environmental conditions^5, 6^. Indeed, a number of previous studies focused specifically on the morphological changes that occur in *B. subtilis* macrocolonies as a result of exposure to antibiotic/antifungal materials, secreted by competing bacterial species such as *Streptomyces*^7^ and *Pseudomonas*^8^.

Chlorhexidine (CHX) is an antiseptic that has previously been shown to have varying antimicrobial effects on a number of biofilm forming bacteria^9^. In *B. subtilis*, it has been shown to damage the cell wall of bacterial cells^10^. On a macro-morphology scale, CHX was shown to disrupt biofilm structure, reducing both bacterial coverage and vitality^11^. In clinical practice, CHX is widely used for dental care in the form of chlorhexidine gluconate mouth wash - so much so that recently concerns have been raised regarding resistance to CHX in oral bacteria^12^.

### Structural features

*B.subtilis’* biofilm robustness has been attributed to, among other features, its complex 3D structure. Specifically, the presence of wrinkled areas within the macrocolonies, in colony type or pellicle form, has been found to be beneficial in terms of improving mechanical resistance and aiding in self-repair^13^. The structure itself is also highly adaptive – wrinkle size and distribution have been found to be dependent on nutrient availability^14^, while also instrumental in facilitating liquid transport throughout *B. subtilis* macrocolonies^15^.

### Cellular composition

*B. subtilis* biofilms are characterized by a complex pattern of cellular differentiation, with different cell types occupying various regions within the biofilm^16^. The resulting cellular diversity has been shown to drive cellular migration^17^ and aid bacterial biofilms in self-healing after cuts to the macrocolony^18^. Cellular composition also determines quorum sensing behavior^19^ as well as interactions with the environment, whether at the biofilm-substrate or biofilm-air interfaces^20^.

### Effectiveness of response to stress conditions

Traditional methods of assessing biofilm robustness often utilize measures that relate to colony response to various types of non-optimal environmental conditions. These include conditions such as mechanical stress^21^, nutrient stress^14, 22^, application of various anti-biofilm agents^23^ and presence of competing species^24^.

Biofilm robustness is thus a multi-parameter characteristic that requires objective tools for biofilm quantification. This is a challenging task, especially given the high phenotypic diversity of *B. subtilis* biofilms, as tools for objective quantification and characterization of undisrupted biofilm macrocolonies are not widely available. Assessing biofilm robustness, however, is necessary to predict potential impact of biofilms, whether in terms of their potential pathogenicity, susceptibility to anti-biofilm agents or mechanical removal.

Several image-based analyses of *B. subtilis* macrocolonies have been published in the literature, with special focus on 3D architecture^25, 26^. In this paper we propose a series of computational analyses of the morphological response of B. *subtilis* biofilms to an antibacterial source. In this model, mature *B. subtilis* colonies were grown on agar substrates with chlorhexidine (CHX) droplets placed in their proximity from the point of seeding. The differences in morphology of the resulting macrocolonies were analyzed computationally and used to establish a now-publicly available software pipeline to analyze mature *B. subtilis* macrocolonies.

## Methods

### Biofilm formation

*B. subtilis* YC161 strain with *P_spank_*-gfp^27^ were grown in Lysogeny broth (LB; 1% tryptone, 0.5% yeast extract, 0.5% NaCl; Neogen, Lansing, MI, USA) and incubated at 37 °C at 150 rpm for five hours. For macrocolony formation, 2.5 μL of starter culture suspension (O.D. 600 nm = 1) was inoculated onto biofilm-promoting LBGM agar medium. LBGM medium was prepared using LB growth medium solidified by the addition of 1.5% (w/v) agar and further supplemented with 1% (v/v) glycerol and 0.1 mM MnS*O*_4_^28^. An amount of 2.5 μL chlorhexidine digluconate (CHX) solution at 20% in water (Sigma-Aldrich, St. Louis, MO, United States) was spotted one-sided at various distances (1.0 cm, 1.5 cm, 2 cm) from the bacterial inoculation point, at the time of inoculation. In addition, control *B. subtilis* biofilm plates were prepared which did not contain CHX. All plates were incubated at 30 °C for a period of three days and images captured daily. Number of repeats were as follows - 12, 11, 9, 12 plates for distances 1, 1.5, 2 cm and control, respectively.

### Fluorescent microscopy

Nikon SMZ25 microscope with ORCA-R2 camera (Nikon Corporation, Tokyo, Japan) was used to capture 2D macrocolony fluorescent images (ND2 format). GFP channel was obtained at excitation/emission of 470/535 nm.

### Computational analysis

PyCharm IDE 2019.2 (JetBrains, Prague, Czech Republic) with Python programming language (Python 3.7, Python Software Foundation, https://www.python.org/), was used for all computational analyses and figure preparation. MATLAB 9.8.0 (R2020a, Natick, Massachusetts, The MathWorks Inc., 2020) script was written for manual selection of colony center coordinates. ND2 files were loaded into 3D numerical matrices of intensity values. Loaded images were converted from 8-bit unsigned integer (Uint-8) into 64-bit floating point.

#### Macrocolony periphery and core segmentations

Captured fluorescent macrocolony images were converted to grayscale, eroded with kernel size [5,5] and then binarized using Otsu’s thresholding method. The resulting binary masks were used as input to OpenCV-Python (cv2) module for contour detection. The contours were then sorted by their descending area coverage. The largest contour (by area) in the output represents the peripheral segmentation of the macrocolonies while the second largest contour represents the colony core.

#### Fit to ellipse

OpenCV-Python (https://pypi.org/project/opencv-python/) module was used to fit contours (periphery and core) to ellipses.

#### Linear regression

Scikit-learn (https://scikit-learn.org/stable/index.html) machine learning library for Python was used to compute a least-squares linear regression.

### Statistical analysis

Scipy^29^ library was used to perform statistical analyses using Student’s t-Test. Statistically significant values were determined by a *p*-value of less than 0.05. Throughout the paper, the symbol * indicates p-value < 0.05, ** indicates *p*-value < 0.01.

## Results

The following section describes a series of morphological measurements of *B. subtilis* macrocolonies’ response to one-sided CHX exposure. The first subsection details changes related to macrocolony growth and expansion, followed by a second subsection which focuses on GFP signal intensity and additional phenomena.

### Expansion kinetics

#### Macrocolony expansion

Figure 1a shows the original macrocolony images, as obtained by fluorescent microscopy 24, 48 and 72 hours after initial seeding. On a macro-scale, Figure 1b demonstrates that there is an inverse relationship between the distance of CHX from the seeding point to the expansion rate of the macrocolony. Macrocolonies that were seeded with CHX at 1 cm (closest) distance exhibited statistically significant reductions in expansion over all three days. In contrast, macrocolonies with CHX at 1.5 cm were smaller in a statistically significant manner only on day 3, while 2 cm macrolonies did not differ from control on any of the days. Table 1 summarizes the relevant *p*-values (two-sided *t*-test).

**Figure 1.**
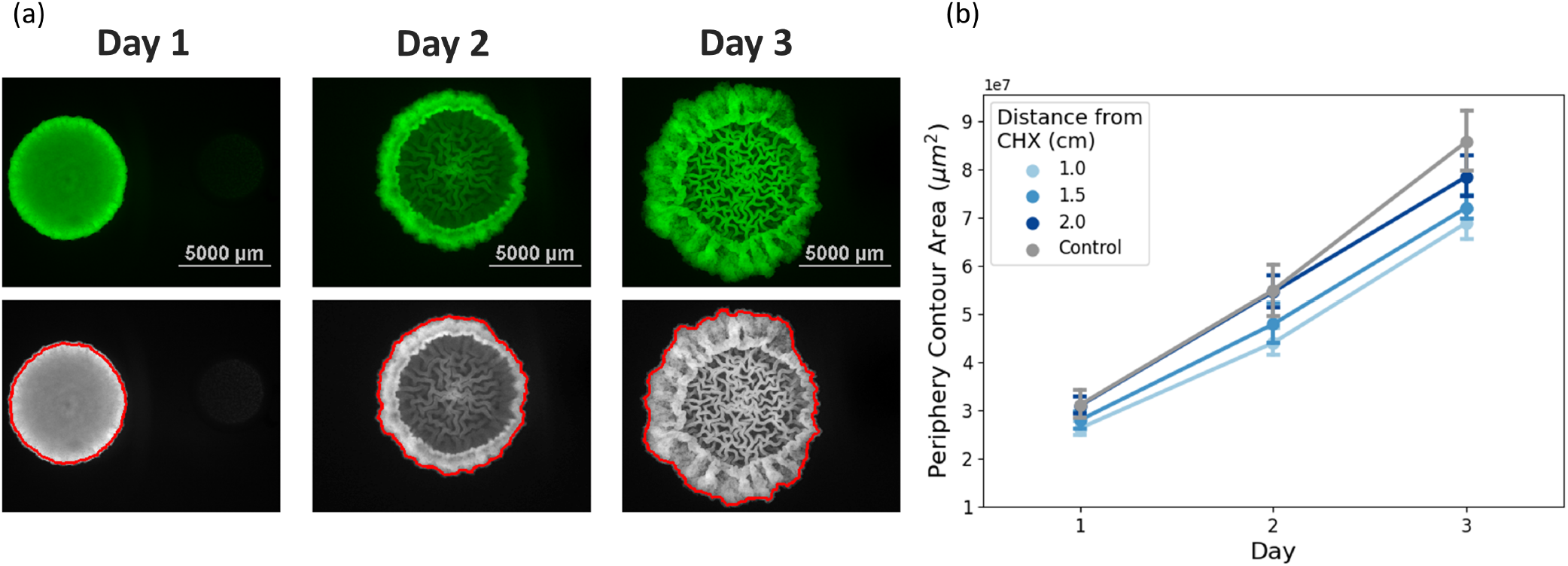
Macrocolony growth and expansion. (a) Fluorescent images of *B. subtilis* macrocolony development over a period of three days. CHX droplet is located horizontally to the right of the macrocolonies in each image, at a distance of 1 cm from the macrocolony center. (b) Total coverage area (μm^2^) of macrocolonies.

**Table 1.** Expansion rates comparison between control and various distances of CHX - *p*-values.

|  | Day 1 | Day 2 | Day 3 |
| --- | --- | --- | --- |
| 1 cm vs. control | 0.01 | 0.002 | 0.0001 |
| 1.5 cm vs. control | 0.07 | 0.05 | 0.001 |
| 2 cm vs. control | 0.44 | 0.85 | 0.1 |

#### Quantifying the morphological response at the periphery

The morphological changes that occur as a result of CHX proximity can be seen on day 2 and 3 – colony periphery on the “exposed” (i.e., right-hand) side of the macrocolony is notably thinner than that on the “unexposed” (i.e., left-hand) side (Figure 1a). In order to quantify the morphological changes that occur in *B. subtilis* macrocolonies as a result of proximity to CHX source during maturation, a series of computational measurements were applied to the images (Figure 2a): firstly, the macrocolony was segmented into an “exposed” and “unexposed” sides by a vertical cut through the macrocolony that directly passes through the colony center (i.e., seeding point) – the separating line is shown in yellow. For each macrocolony, a binary image was obtained using Otsu’s thresholding method. For each macrocolony, an outer contour surrounding the entire macrocolony was determined using a border following algorithm applied on binary images from the previous step – the resulting contour is shown in red. For both the exposed and unexposed sides, a half-contour was mirrored around the separating line. The resulting mirrored contours can be seen in Figure 2a, middle column – top image shows the unexposed side contour, as it was mirrored onto the exposed side, while the bottom image does the same for the contour of the exposed side. Each one of the two contours is then fitted to an ellipse, shown in white – Figure 2a, rightmost column. The semi-major and semi-minor axes of the fitted ellipses were measured.

**Figure 2.**
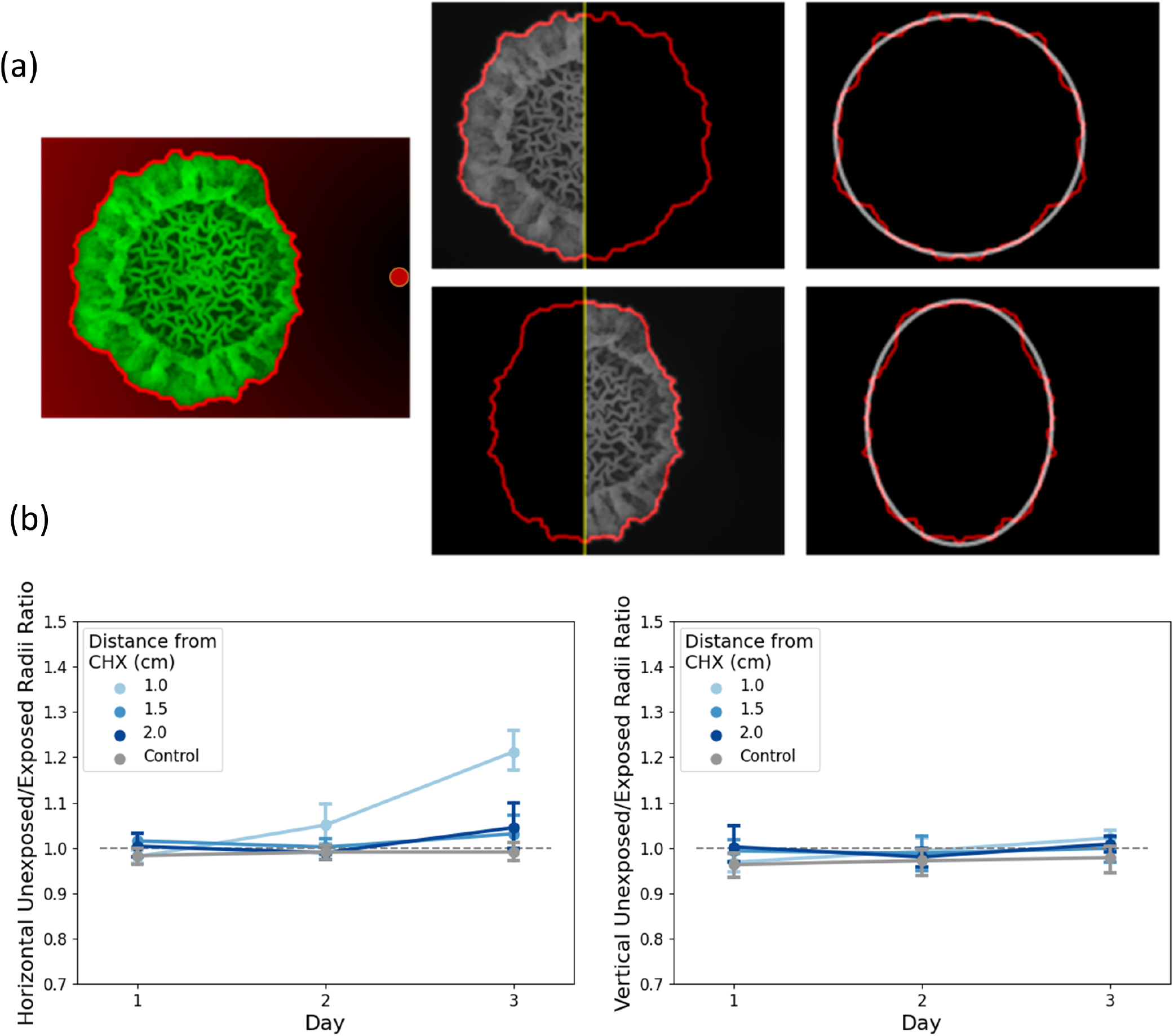
Illustration of inhibition measurement at the periphery. (a) The macrocolony is divided vertically into unexposed (left) and exposed (right) halves. The CHX spot is horizontal to the right of the macrocolony in each image. Each macrocolony half is separately mirrored and the resulting contour fitted to an ellipse. Red background in leftmost image reflects the Euclidean distance of each pixel from the CHX source. Outer contours are shown in bright red. (b) **Colony periphery deformation analysis.** At each distance from CHX source (control and 1/1.5/2 cm) the ratio between horizontal (left) and vertical (right) radii between the unexposed and exposed halves is shown.

Figure 2b demonstrates the differences in morphology that occur between the exposed and unexposed sides, both in the horizontal (left) and vertical (right) planes. The loss of symmetry that occurs in macrocolonies as a result of CHX proximity on day 3 is statistically significant in the horizontal and vertical planes only in macrocolonies with CHX placed at a distance of 1 cm. Thus, changes in morphology are directly correlated to the distance from the CHX source.Table 2 summarizes the relevant *p*-values (two-sided *t*-test).

**Table 2.** Horizontal and vertical inhibition rates comparison between control and various distances of CHX - *p*-values.

|  | Day 3 - horizontal | Day 3 - vertical |
| --- | --- | --- |
| **1 cm vs. control** | *< 0.0001* | *0.03* |
| **1.5 cm vs. control** | 0.1 | 0.33 |
| **2 cm vs. control** | 0.06 | 0.13 |

### Quantifying the morphological response at the core

Figure 3a illustrates the same image processing pipeline, applied to the colony core, rather than the periphery. Figure 3b demonstrates that no comparable changes in morphology occur at the colony core, whether in the horizontal (left) or vertical (right) planes. Indeed, no statistically significant loss of symmetry was observed at the colony core, regardless of distance from CHX source.

**Figure 3.**
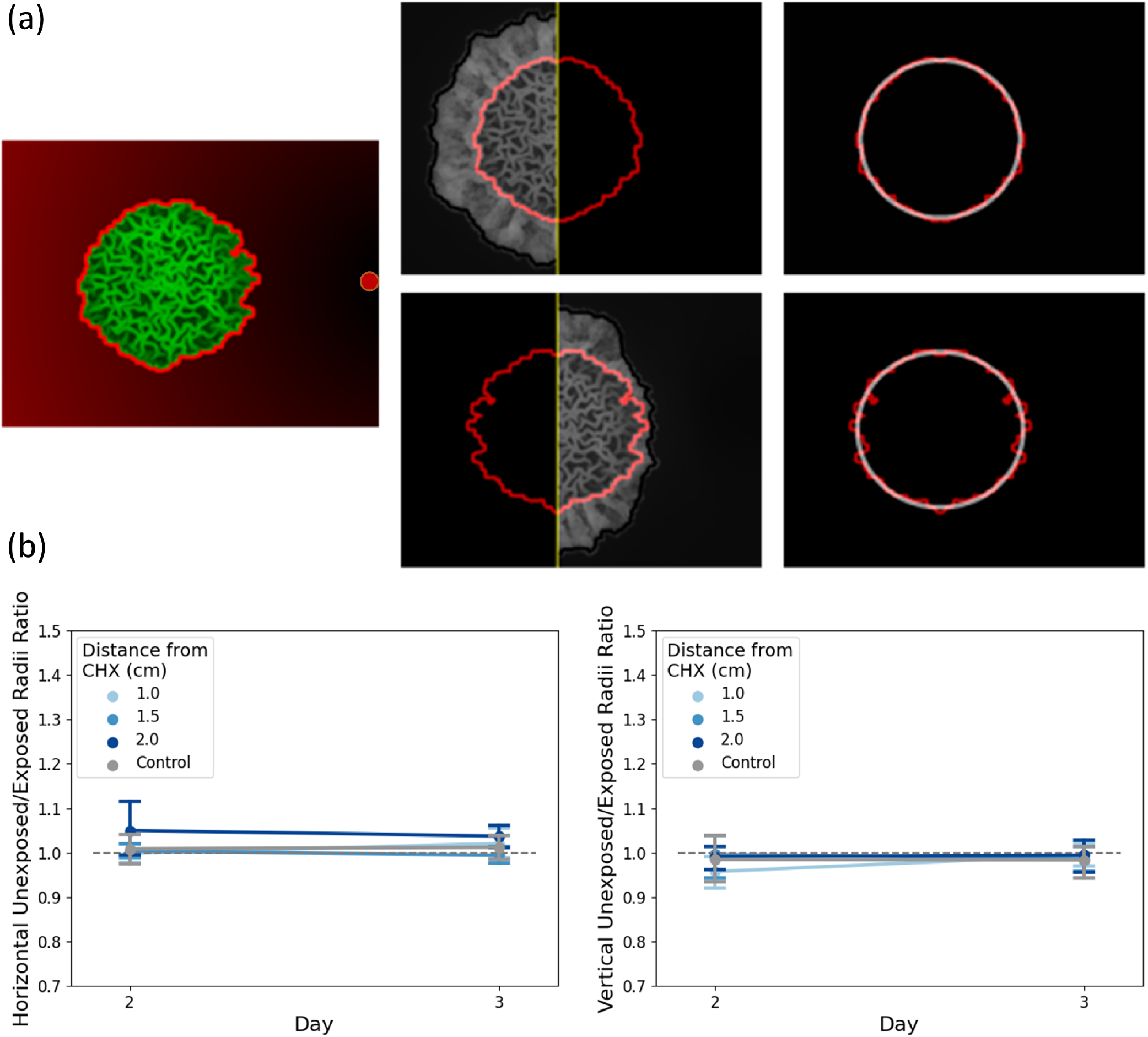
Illustration of inhibition measurement at the core. (a) Illustration of inner core segmentation with mirroring and fitting to ellipse. (b) **Colony core deformation analysis.** At each distance from CHX source (control and 1/1.5/2 cm) the ratio between horizontal (left) and vertical (right) radii between the unexposed and exposed halves is shown.

Figure 3b shows that on day 3, macrocolony core did not differ in a statistically significant manner from the control, regardless of CHX proximity. The colony core is therefore more preserved in structure than colony periphery (or more resistant to CHX). Table 3 summarizes the relevant *p*-values (two-sided *t*-test).

**Table 3.**
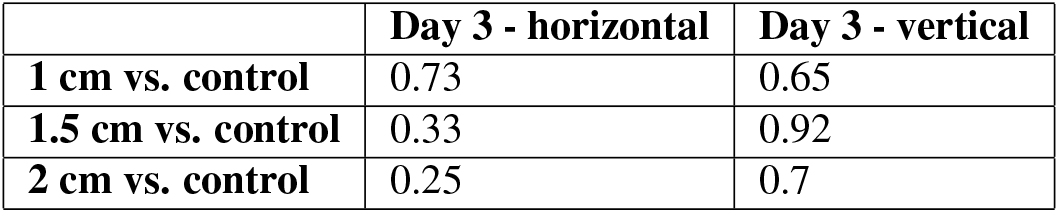
Horizontal and vertical inhibition rates comparison between control and various distances of CHX - *p*-values.

### Intensity analysis

#### Effect of proximity to CHX on bacterial viability

Figure 4a illustrates how distance from CHX is determined for each pixel in the macrocolony. Euclidean distance was used for the calculations. Figure 4b demonstrates how pixel intensity is affected by proximity to CHX source: average pixel intensity at the exposed/control areas is shown for both periphery (orange) and core (blue) regions on day 3. In other words, for each macrocolony, the ratio between average pixel intensity of the exposed and unexposed halves was calculated and compared at the periphery and core regions. Statistically significant differences in values were found in periphery of macrocolonies that were grown at distance 1 cm from CHX, as well as core of macrocolonies that were grown at distance 1.5 cm from CHX. Thus, at these distances, the macrocolony is affected both by morphological deformation as well as changes in GFP intensity.

**Figure 4.**
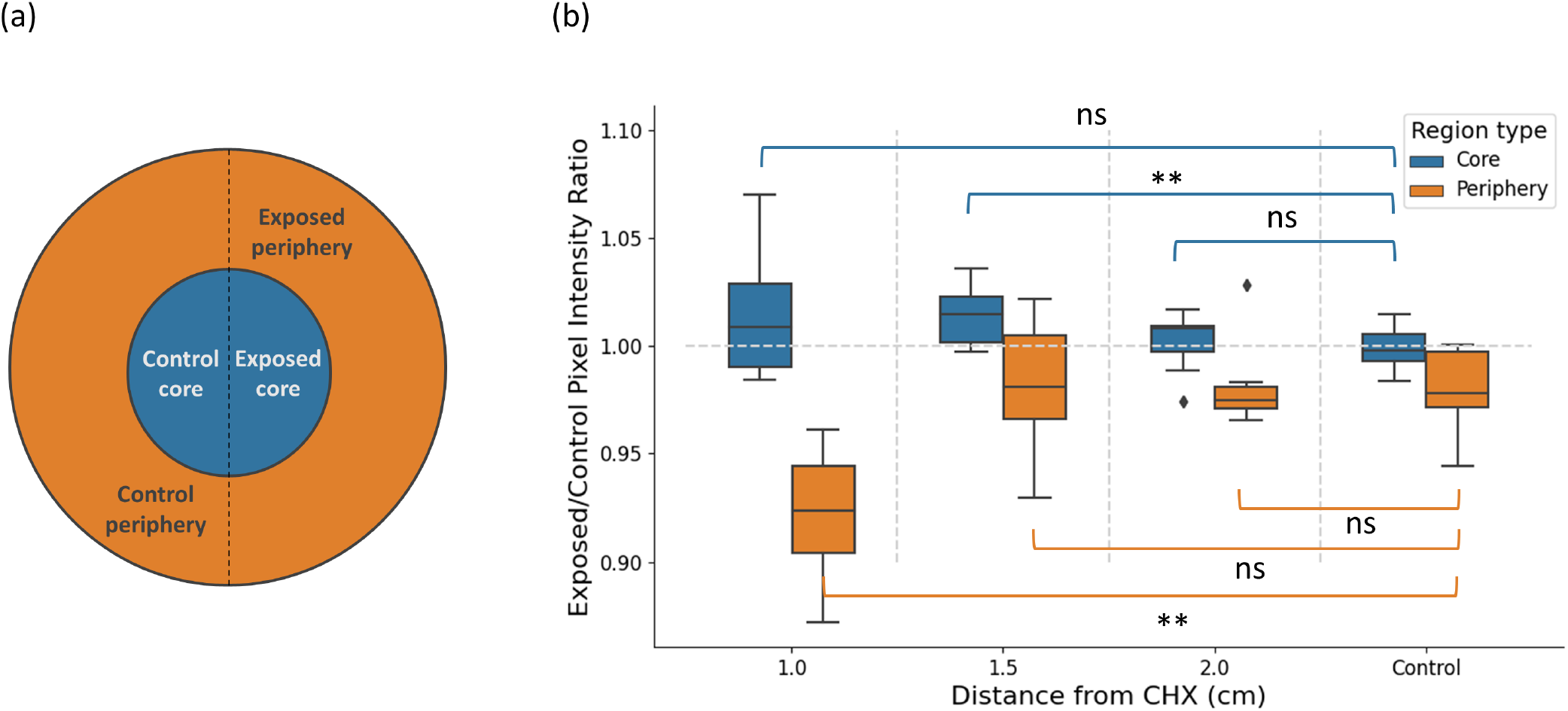
Pixel intensity calculation. (a) Image illustrating the different areas within the macrocolony. CHX source lies directly horizontally to the right. (b) Ratio of intensity average between unexposed and exposed sides of the macrocolony is shown separately for the periphery (orange) and the core (blue). For control images, unexposed and exposed sides were determined via data augmentation as average ratio of left vs. right halves, top vs. bottom halves and a combination of upper left and bottom right quadrants vs. upper right and bottom left quadrants. The shorthand “ns” indicates non-significant *p*-value (> 0.05).

#### Correlation between GFP intensity and distance to CHX

Bacterial cells at the leading edge of the macrocolonies are those that are located at the outermost layer of the macrocolony periphery. Due to the curvature of the macrocolony, points along the leading edge are located at varying distances from the CHX source (Figure 5a). In order to characterize the nature of relationship between pixel intensity and distance to CHX, pixel intensities along the leading edge were plotted in Figure 5: red dots represent pixels along the leading edge of the exposed side of a macrocolony grown at 1 cm from CHX, while blue dots represent pixels along the leading edge of the exposed side of a macrocolony grown at 2 cm from CHX. Given both sets of pixel intensity values, a linear regression model was applied to both – as can be seen in Figure 5, there is a linear correlative relationship between Euclidean distances and pixel intensities. This relationship is stronger when CHX is located closer to the macrocolony center – for example, in the images that are shown in Figure 5, linear approximation revealed that 1 cm macrocolonies are characterized by a slope that is over three times higher (red) than that of the 2 cm macrocolonies (blue). This finding signifies the linear relationship between GFP signal intensity of cells located at the leading edge of the macrocolony to their distance from the CHX source.

**Figure 5.**
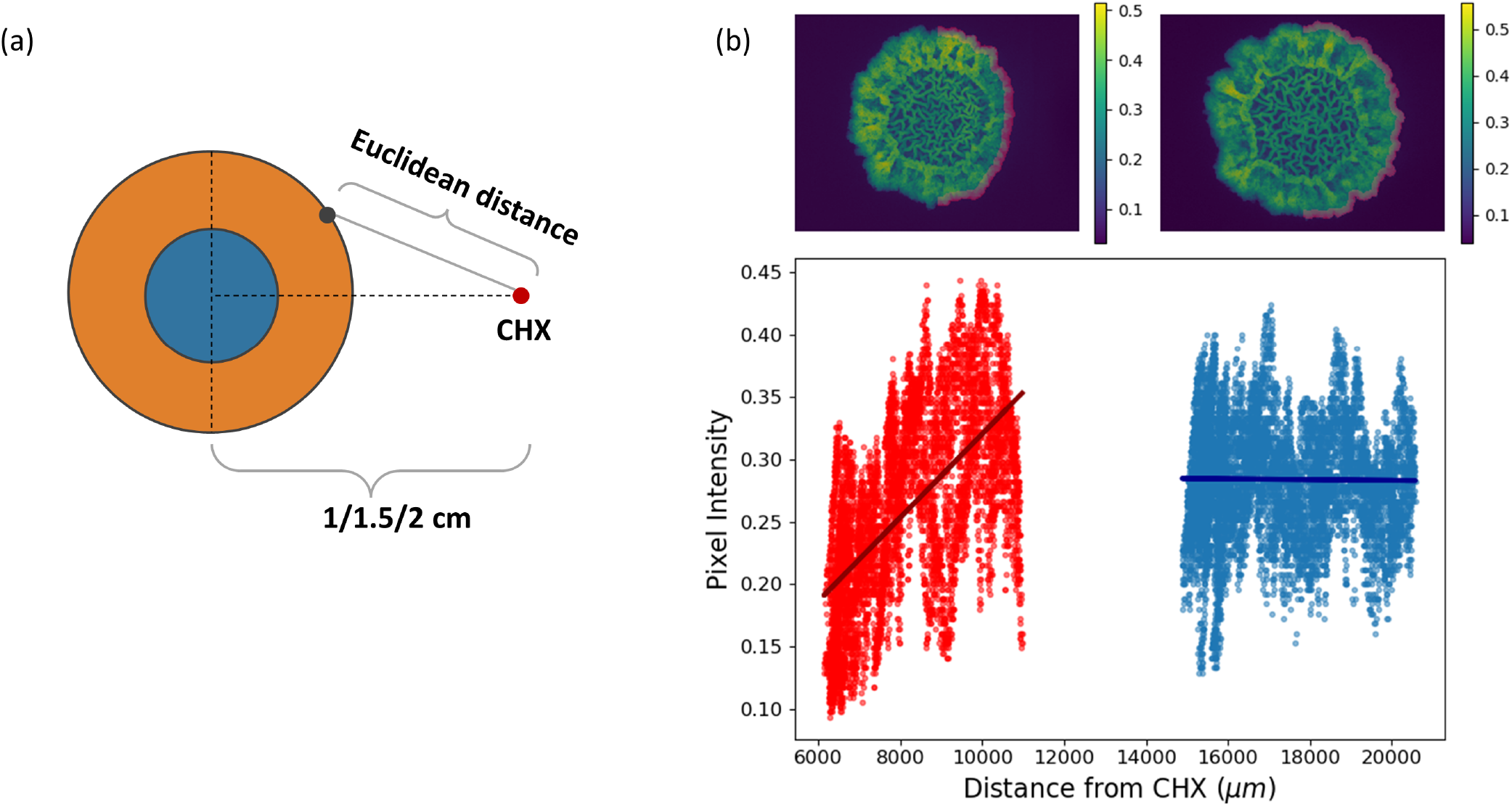
Linear regression model for pixel intensity at the leading edge as function of Euclidean distance from CHX source. (a) Illustration demonstrating the distance calculation between the CHX source (red dot) to each pixel within the macrocolony. (b) (top) *B. subtilis* macrocolonies on day 3 at 1 cm (left) and 2 cm (right) distances from CHX source. (bottom) Intensities of pixels located at the leading edge (highlighted 20 pixels-wide section from the outer rim) of the exposed half of the macrocolony: red pixels originate from 1 cm macrocolony, blue pixels originate from 2 cm macrocolony. Linear regression lines demonstrate that at 1 cm, pixel intensity is correlated to the distance from CHX source, while no such effect is seen at 2 cm macrocolony.

### Crescent shaped morphology

As CHX is placed closer to the macrocolony, it exerts greater inhibitory effect, resulting in increasing deformation of the macrocolony on the side closer to the antimicrobial source. However, in the case when CHX is placed at 0.5 cm from the initial point of seeding the macrocolony only develops towards the unexposed side – Figure 6a shows the growth of a sample macrocolony over a period of three days (left-to-right). Starting from day 1, the macrocolony appears to grow only on the side opposite CHX location. On average, control macrocolonies expand on day 1 to a radius of 0.3 cm. CHX droplet is on average 0.2 cm in radius. Hence, even when CHX is placed at a distance of 0.5 cm, the macrocolonies have enough potential space to expand to 0.3 cm. However, Figure 6d demonstrates that despite the fact that there is sufficient unoccupied space in front of the macrocolony to expand into (indeed, equal to that required by control macrocolonies which are uninhibited by CHX), the macrocolony does not expand towards the exposed side at all. Rather, it expands towards the opposite side and consequently assumes a unique ‘crescent’ shape starting from day 1 onwards.

**Figure 6.**
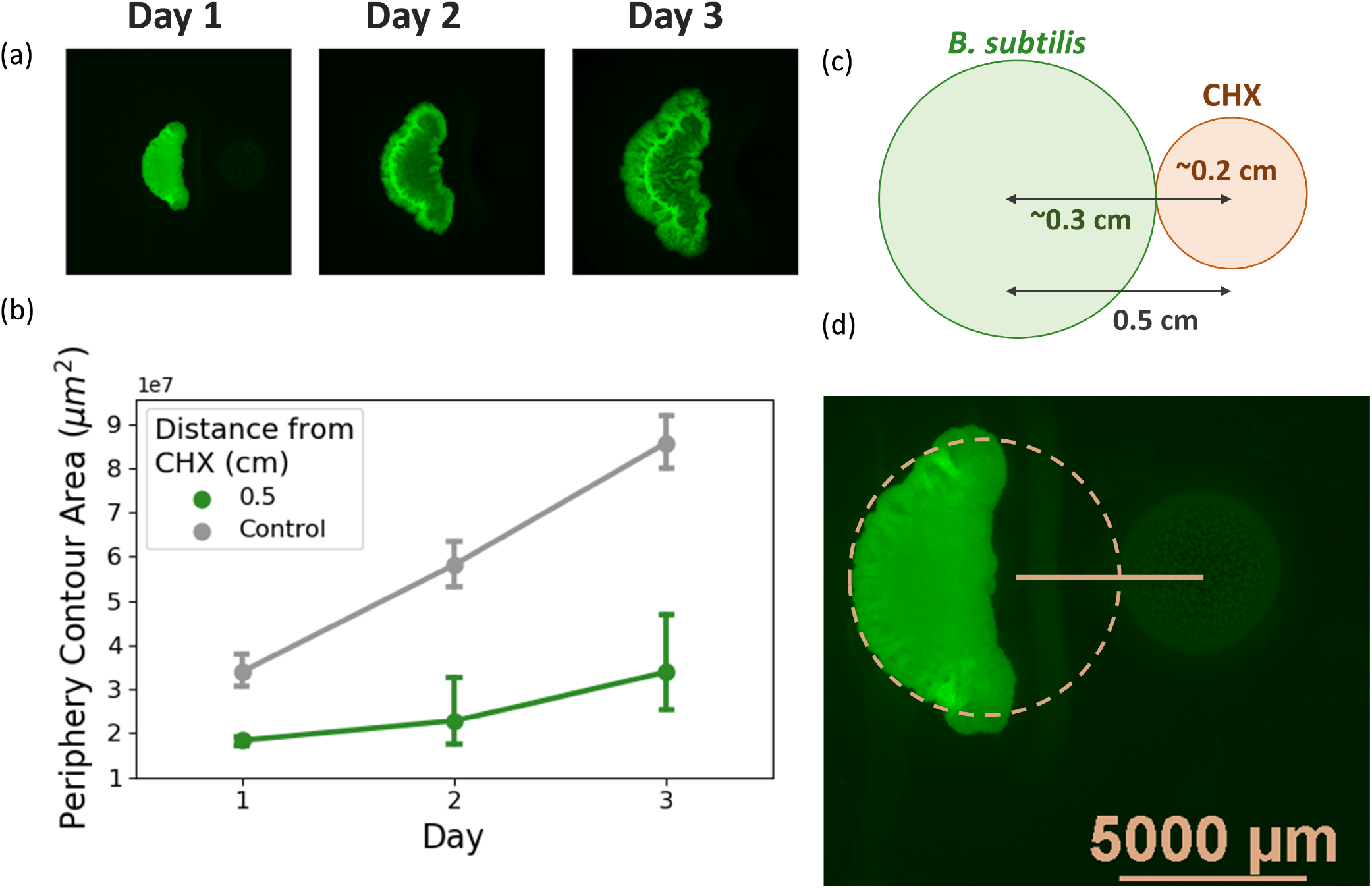
Crescent shaped morphology occurring at CHX distances of 0.5 cm. (a) Top row illustrates the macrocolony morphology over a period of three days – the change in morphology appears in the form of “crescent shaped” colonies. (b) Expansion comparison with control macrocolonies. (c) Illustration depicting several relevant distances when CHX is placed at 0.5 cm – average radius of a mature B. *subtilis* macrocolony on day 1, average radius of a CHX droplet. (d) Representative image of macrocolony on day 1, with CHX (color corrected for visual clarity) shown to the right.

### Effects seen in the agar substrate

Figure 7a shows bright field images of B. *subtilis* macrocolonies, with CHX droplets seen to their right. This visualization reveals a bright formation in the agar substrate, between macrocolony and CHX, undetected in the fluorescent images. This structure is embedded into the agar throughout its entire width, as seen in Figure 7b. Over a period of three days, its shape changes from concave to convex, with it seemingly ‘engulfing’ the CHX droplet. More interestingly, the appearance of the agar at both sides of the formation is uneven, best visualized in Figure 7b, where the agar on the CHX side appears “muddy”, unlike the one on the macrocolony side.

**Figure 7.**
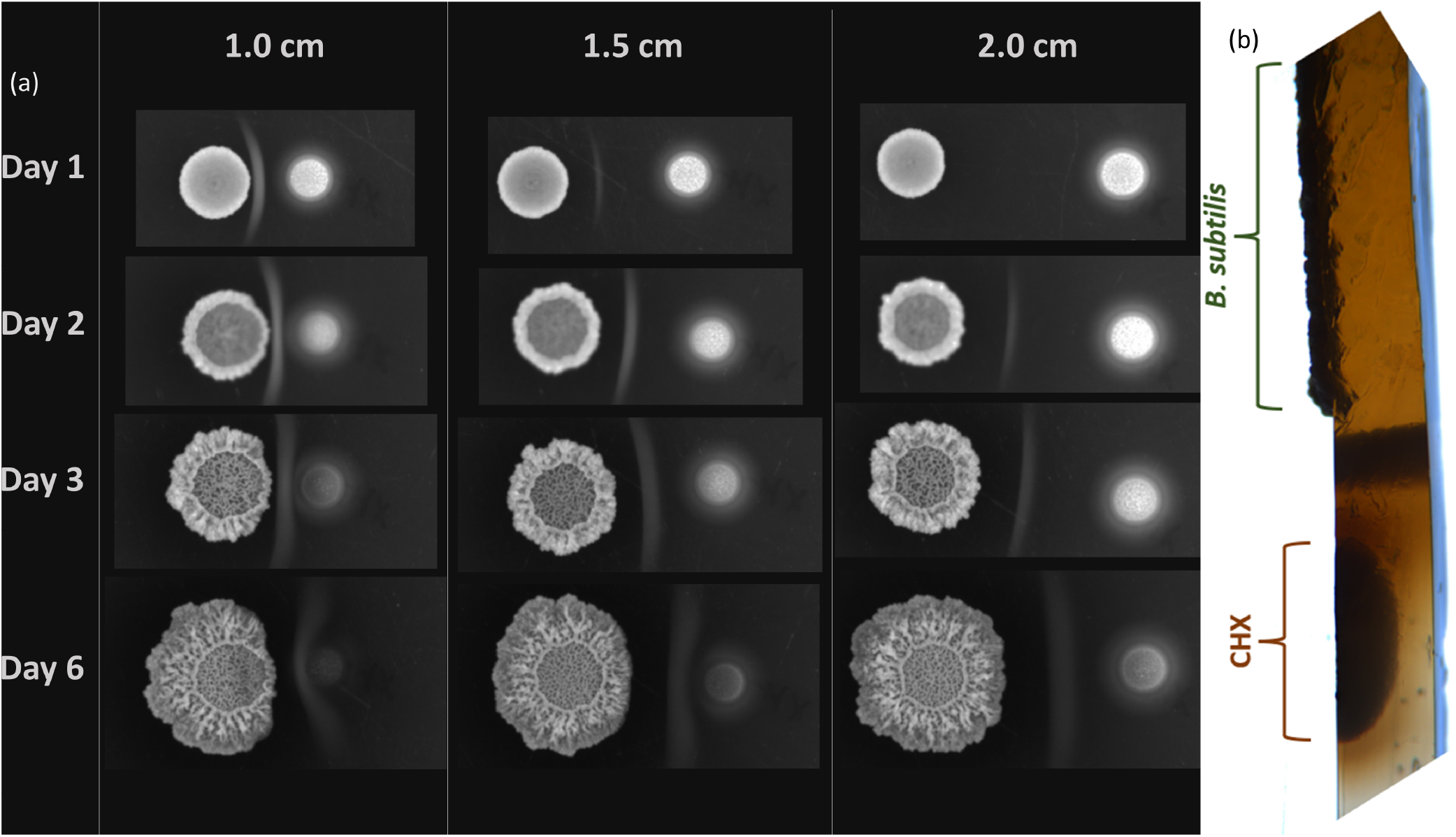
Bright field images of expanding macrocolonies. (a) Bright field images of expanding *B. subtilis* macrocolonies grown in proximity to CHX at 1/1.5/2 cm. CHX droplet is seen to the right of the macrocolonies. (b) Cross-section of agar substrate seeded with a macrocolony and CHX droplet.

## Discussion

### Core vs. periphery

The subdivision of biofilm macrocolonies into “core” and “periphery” regions is characteristic of *B. subtilis* biofilms, but it is not unique to them. Indeed, a number of bacterial species exhibit similar biofilm morphology including the pathogens *Escherichia coli*^30, 31^ and *Staphylococcus aureus*^32^. This phenomenon puts forward a question of whether these regions differ in their respective roles in the life of biofilms. In order to answer this question, it is necessary to develop tools for accurate quantification of various features of these two regions. In this context, computational assays performed on images of biofilm macrocolonies provide a dual benefit - they are both reproducible and allow for quantifiable comparison of samples - e.g., biofilms treated with different agents.

In the literature, several differences have been established between the ‘core’ and ‘periphery’ regions of *B. subtilis* biofilms. Specifically, the core region, often recognized by a complex mesh of ‘wrinkled’ structures in mature biofilms, was found to be characterized by reduced expansion rates and reduced long-term cellular viability, compared to the periphery^33^. The role of the wrinkled structures has been suggested to aid in solute transport^15^, nutrient uptake^14^ and self-repair^18^.

This paper demonstrates that the core region develops more symmetrically in proximity to CHX than the periphery. In other words, it seems that its growth on the side exposed to CHX is less affected, compared to the periphery. Indeed, while periphery regions of the exposed and control sides are significantly different in both area occupied (Figures 2, 3) and GFP signal intensity (Figure 4), corresponding core regions do not exhibit comparable differences.

Possible causes for this difference in region ‘robustness’ may lie in molecular properties – cellular content and extracellular matrix composition. For example, calcium precipitates were previously identified at wrinkled regions in *B. subtilis* biofilms^34^. Calcium was previously found to be a stabilizing agent in *B. subtilis* biofilm structure^35^ and its presence at the wrinkled regions of the colony core^34^ may serve as a physical barrier that resists deformation of the core region, effectively “walling off” the core. *B. subtilis* macrocolony responses to proximity to other types of bacteria has been studied in the literature – for example, in response to *Staphylococcus epidermidis*, the colony was shown to expand towards the offending source, rather than recede from it^36^. In response to its close relative *B. simplex, B. subtilis* macrocolonies engulf and envelop its macrocolonies – expression of motility genes is required for this to occur^37^. Thus, *B. subtilis* employs a number of strategies, depending on the source.

The changes seen in the substrate may be the result of one of the strategies that the bacterium employs to effectively distance itself from an offending source – it is likely that both mechanical and molecular machinery is involved. Under fluorescent microscopy, the formation does not emit a GFP signal and therefore is unlikely to contain live bacterial cells. The structure is distinct from the brown pigment secreted into the agar by mature *B. subtilis* macrocolonies and is possibly a result of CHX precipitation with various proteins secreted by the macrocolony. Further studies may shed light on the composition of the agar at the interface between macrocolonies and CHX, possibly revealing a new mechanism for biofilm ‘robustness’.

### Intensity correlation to distance from CHX

GFP signal intensity along the leading edge of *B. subtilis* macrocolonies was found to be correlated to Euclidean distance from CHX. However, Figure 5 demonstrates that while such correlation exists, at each distinct Euclidean distance, the distribution of GFP values of pixels at that distance is extensive. This can be seen by the relatively wide distribution of pixel intensities along the Y-axis, for each value on the X-axis. As can be seen in Figure 5b, on average, for each distance value, pixel intensities range is approximately 0.2 points wide, in both 1 cm and 2 cm samples. This suggests a natural variation in cell density (as reflected by GFP signal intensity) along the leading edge of a macrocolony. While this variation remains preserved in both samples regardless of distance from CHX, in macrocolony where CHX was placed at 1 cm, there exists a correlation between average GFP intensity and Euclidean distance from CHX. Thus, a distance-based effect exists where areas closest to CHX are less dense than those located farther from it.

### Loss of core symmetry

With decreasing distances of CHX from the colony center, its effect on morphology is enhanced – on a macro scale, the macrocolony is characterized by reduced expansion at the side exposed to the CHX source. At CHX distances between 1 and 2 cm, colony core remains preserved in symmetry (Figure 2b). However, at a distance of 0.5 cm the macrocolony seems to ‘abandon’ growth on the exposed side, but rather grow as a ‘crescent-shaped’ macrocolony from day 1 (Figure 6). Such morphology is distinct from those observed at larger CHX distances – CHX droplet radius, which is measured as approximately 0.2 cm, still leaves enough space for the macrocolony to expand. As the average radius of control macrocolonies was measured as approximately 0.3 cm, there is enough space to expand at least until day 1. However, not only is there zero expansion of the exposed side, the macrocolony core seems to be receded even onto the unexposed side. This creates the ‘crescent-shaped’ macrocolonies with concave boundary facing the CHX droplet. Nonetheless, the complex wrinkled core remains intact, as it develops at the center of what would have been the control (unexposed) side of the macrocolony. This phenomenon suggests that the change in overall macrocolony morphology (into crescent shape) occurs at CHX distances smaller than 1 cm, but greater than 0.5 cm. Future studies may not only determine this distance value, but use it as another numerical parameter of biofilm robustness.

## Conclusion

Computational analyses such as those presented in this paper offer a new method for analyzing biofilms in terms of the morphological changes that occur in response to antimicrobial agents. Indeed, our computational framework supports analysis of different inhibitors such as antibiotics, antifungals and more. Given parameters that were measured in this paper, one can compare them directly between different macrocolonies, e.g., those grown under varying nutrient availability. Do macrocolonies grown under non-optimal nutrient conditions retain their ability to respond to CHX proximity with the same effectiveness as macrocolonies grown under optimal nutrient conditions? In other words, are such macrocolonies able to retain the symmetry of the core region at comparative CHX distances/concentrations? It is our hope that this paper, along with its publicly available code, will make is possible to further study such biofilm properties and bring us closer to a comprehensive model of comparative biofilm ‘robustness’.

## Data availability

All code and datasets relevant to this manuscript can be found at - https://github.com/cohenoa/An-Open-Source-Computational-Tool-for-Measuring-Bacterial-Biofilm Installations and program execution instructions are available both in the code repository as well as a supplementary file to this paper.

## Acknowledgements

This research was partially supported by the STEP Graduate Training Program (STEP-GTP), the Dr. Izador I. Cabakoff Research Endowment Fund and the Azrieli College of Engineering Research Fund.

## Author contributions statement

S.G., N.E.C. and D.S. conceived the idea. S.G. and R.S. designed and performed the biological experiments. S.G. and N.E.C. designed and performed the computational analyses. S.G., N.E.C. and D.S. wrote the paper. R.S. and O.F. revised the manuscript critically and provided academic guidance. All authors have read and agreed to the published version of the manuscript.

## Additional information

The authors declare no competing interests.

## Notes

https://github.com/cohenoa/An-Open-Source-Computational-Tool-for-Measuring-Bacterial-Biofilm-Morphology-and-Growth-Kinetics

### Competing Interest Statement

The authors have declared no competing interest.

### Summary of Updates

Title changed

